# Quantitative 3D imaging of intact engineered heart tissues using inverted selective plane illumination microscopy

**DOI:** 10.64898/2026.10.08.757603

**Authors:** Kaisla Walls, Lotta Kulmala, Joose Kreutzer, Antti Ahola, Mari Pekkanen-Mattila, Jari Hyttinen, Birhanu Belay

## Abstract

**Background:** Advanced *in vitro* models of the human heart have become promising tools for studying cardiac diseases and drug discovery. Three-dimensional (3D) engineered heart tissues (EHTs) provide more adult cardiac tissue-like structural organization compared to traditional two-dimensional (2D) *in vitro* models with straightforward contractile readouts. However, the assessment of the architecture of EHTs, especially the quantification of cellular and tissue structural components, has remained poorly studied. Here we demonstrate the use of light-sheet microscopy for structural imaging of intact EHTs and provide quantification of the EHT cellular and subcellular structures.

**Methods:** We developed an in-house inverted selective plane illumination microscopy (iSPIM) platform and combined it with iterative indirect immunofluorescence (IT-IF) staining for better signal acquisition. We developed an image-processing and analysis pipeline dedicated to volumetric EHT analysis.

**Results:** With this integrated platform, we achieved volumetric imaging of intact EHTs and quantification of 3D tissue architecture, revealing aligned cardiomyocytes within the imaged volume and increasing cardiac nuclei content towards the core of the tissue. The developed methods allowed the detection of heterogeneities between tissue sections, demonstrating the sensitivity required for future disease modeling or maturation studies.

**Conclusions:** The developed integrated platform opens new possibilities for volumetric analysis and characterization of 3D tissue structure using iSPIM, providing new insights into EHT architecture and the use of EHTs as disease modeling tools.

## 1. Introduction

Cardiovascular diseases (CVDs) remain the leading cause of mortality worldwide [1,2], highlighting the need for physiologically relevant experimental models. Human-induced pluripotent stem cell (hiPSC) technology has enabled the development of cardiac *in vitro* models that recapitulate patient-specific genetic background [3,4]. However, two-dimensional (2D) cell cultures fail to capture the three-dimensional (3D) structural organization and function inherent to human myocardium. To address these limitations, engineered heart tissues (EHTs) have emerged as advanced 3D cardiac models that provide more physiologically relevant *in vitro* modeling [5]. EHTs are increasingly used for studies of disease modeling and drug testing [6–8], while also enabling direct assessment of force generation [9,10] and action potentials [11,12]. Yet, characterization of the structural organization of intact EHTs remains limited, particularly at the whole tissue level. This limitation is important because myocardial structure strongly influences function [13], and is markedly altered in many CVDs, including myocardial infarction [14] and hypertrophic cardiomyopathy [15]. In this study, we developed an imaging and analysis workflow for visualization and quantitative assessment of intact EHT structure in 3D.

EHTs undergo substantial structural remodeling during culture and maturation [5], with increased cellular alignment, a higher cytoplasm-to-nucleus ratio, and an increased force generation [16]. At cellular level, hiPSC-derived cardiomyocytes (CMs) exhibit a more adult-like morphology when cultured in EHTs with length-to-width ratio of 4:1 [16], compared to more round shape found in 2D cultures [17]. Similarly, expression of sarcomeric and calcium handling proteins is enhanced in EHTs compared with 2D cultures, although substantial differences from human myocardium remain [5]. In heterotypic EHTs, this maturation process also involves spatial organization of different cell populations from endothelial cells (ECs), cardiac fibroblasts (CFs), and smooth muscle cells (SMCs), resulting in clear layering with ECs residing around the cardiac tissue and forming an in vivo-like interface [18]. Additionally, heterotypic EHTs show increased levels of gap-junctional protein connexin 43 (Cx43), an important subunit for electrical coupling structures in cell-to-cell connections, which could contribute to the detected increase in conduction velocity within the EHT [18]. Tissue remodeling has been demonstrated in engineered cardiac models of myocardial infarction, where combination therapies yielded greater recovery of tissue architecture and functional performance than single-treatment approaches [19]. Therefore, both tissue maturation and disease modeling necessitate tools capable of quantifying cellular heterogeneity and spatial organization to resolve structural changes and cellular relationships underlying tissue function, remodeling, and disease.

Structural analysis of EHTs remains technically challenging as these constructs can range from several hundred micrometers to millimeters in thickness [18,24], resulting in increased optical scattering and reduced imaging depth. The two-pole EHTs are formed and cultured between poles that influence tissue organization and function through passive mechanical loading [24]. Preserving native geometry of the tissue during imaging is therefore important for accurate structural assessment. However, many commonly used imaging and histological methods require removal of the tissue from the poles or physical sectioning, which can alter tissue geometry and compromise the analysis of EHTs in their native functional state. This is particularly important because structural features are closely linked to cardiac tissue function and maturation, and may vary throughout the tissue volume [25]. Consequently, analyses based on isolated sections or projected images may not accurately represent the 3D architecture of the entire tissue. Conventional fluorescence imaging methods, such as confocal microscopy, are often limited to analysis of relatively thin tissue sections due to restricted imaging depth, photobleaching, and slow volumetric acquisition [26,27]. Although optical clearing can improve imaging depth in thick tissues, clearing protocols typically involve extensive sample processing and may alter tissue structure [28]. Advanced imaging approaches are therefore needed to enable non-destructive, volumetric, and physiologically relevant characterization of EHT structure in situ.

Selective plane illumination microscopy (SPIM) enables rapid volumetric imaging of large biological samples. By illuminating only the imaging plane and detecting fluorescence orthogonally, SPIM provides optical sectioning with reduced photobleaching and phototoxicity [29]. These capabilities have led to the use of SPIM from whole embryo imaging to visualization of cellular dynamics in 3D [30–33]. Continued technological development has further expanded SPIM capabilities through advances in light-sheet generation [30,34,35], image acquisition speed [36–38], and isotropic-resolution imaging [36,39,40]. In addition, multiple SPIM configurations have been developed to address diverse sample geometries, preparation protocols, and imaging requirements [41]. Among these configurations, inverted selective plane illumination microscopy (iSPIM) allows imaging of samples prepared on standard microscopy substrates and is compatible with conventional sample handling workflows [42]. These characteristics make iSPIM setup well-suited for imaging intact EHTs in their native cultured configuration, although such applications have not previously been demonstrated. Furthermore, the quantitative potential of volumetric imaging remains largely unrealized for EHT analysis.

In this study, we sought to establish a workflow for volumetric structural characterization of intact EHTs by integrating a developed iSPIM platform, iterative indirect immunofluorescence (IT-IF) staining and quantitative image analysis. We first assessed the optical characteristics of the iSPIM system and image processing pipeline using fluorescent beads and zebrafish samples. Additionally, we optimized fixed EHT preparation for enhanced fluorescence imaging. We then applied the workflow to intact EHTs labeled for cardiac-specific troponin T (cTnT), gap-junctional marker connexin 43 (Cx43), and nuclei to quantify nuclei distribution and CM alignment throughout tissue volume. By providing depth-resolved structural information from intact tissues, this approach could support future efforts to validate engineered tissue models and assess disease-associated or pharmacologically induced tissue remodeling.

## 2. Materials and Methods

### 2.1 Ethical approval

Experiments involving zebrafish (*Danio rerio*) embryos were performed at stages not regulated by EU Directive 2010/63/EU (≤5 days post fertilization, dpf), and thus no project specific ethical approval was required. The care, husbandry, and breeding of adult zebrafish were performed in the authorized animal facility of Tampere University, operating under approval ESAVI/44000/2025. The establishment, culture, and differentiation of hiPSC lines were conducted in accordance with the approval of the Ethics Committee of Pirkanmaa Hospital District (Aalto-Setälä R08070). All experiments were performed in accordance with relevant guidelines and regulations.

### 2.2 Differentiation of human induced pluripotent stem cell derived cardiomyocytes

The EHTs were constructed from hiPSC-derived CMs. The previously established cell line UTA.04602.WT, derived from a healthy 55-year-old female, was utilized in this study [43]. For culturing, the hiPSCs were maintained on mouse embryonic fibroblast feeder cells (MEF, Gibco, ThermoFisher Scientific, USA) in KnockOut Serum Replacement (KSR) medium (Gibco, ThermoFisher Scientific, USA), supplemented with 1% Minimum Essential Medium Non-Essential Amino Acids (MEM NEAA, Gibco, ThermoFisher Scientific, USA), 1% GlutaMAX (Gibco, ThermoFisher Scientific, USA), 0.2% β-mercaptoethanol (Gibco, ThermoFisher Scientific, USA), and 0.5% penicillin/streptomycin (Euroclone SpA Milan, Italy).

The hiPSC-CMs were generated following a published embryoid body (EB) differentiation protocol [44]. On day 0, hiPSCs were detached using Versene (Gibco, ThermoFisher Scientific, USA) resuspended in KSR medium supplemented with 5 μM blebbistatin (Sigma-Aldrich, USA). The cell suspension was then transferred to ultra-low attachment 6-well plates to facilitate EB formation. On day 1, the medium was replaced with RPMI 1640 + GlutaMAX (Gibco, ThermoFisher Scientific, USA) supplemented with 1% B-27 Minus Insulin (50X, Gibco, ThermoFisher Scientific, USA), 0.5% penicillin/streptomycin, 5 μg/ml ascorbic acid (Sigma-Aldrich, USA), 10 ng/ml bone morphogenic protein 4 (BMP4, R&D Systems, USA), and 25 ng/ml Activin A (Peprotech, ThermoFisher Scientific, USA). On day 3, the cells received fresh differentiation medium supplemented with 5 μg/ml ascorbic acid. On day 4, the medium was exchanged for differentiation medium containing 2.5 μM IWP-4 (Tocris, Bio-Techne, Ireland) and 5 μg/ml ascorbic acid.

On day 7, half of the medium was replaced with differentiation medium supplemented with 5 μg/ml ascorbic acid. From day 9 onward, half-medium changes were performed three times per week with RPMI 1640 + GlutaMAX supplemented with 1% B-27 Minus Insulin. Beginning on day 11, RPMI 1640 + GlutaMAX was supplemented with B-27 Supplement (50X, serum-free, Gibco, ThermoFisher Scientific, USA).

Following differentiation, hiPSC-CMs were purified by magnetic-activated cell sorting (MACS) using the PSC-Derived Cardiomyocyte Isolation Kit (Miltenyi Biotec), according to the manufacturer’s instructions. The purified hiPSC-CMs were subsequently cryopreserved in CryoStor cell cryopreservation media (Sigma-Aldrich, USA) and stored at −80 °C until use.

### 2.3 Engineered heart tissue preparation

The two-pole EHTs were constructed following a previously published protocol [24], modified to reduce reagent consumption by decreasing the number of hiPSC-CMs to 300,000 cells per EHT. Cryopreserved hiPSC-CMs were thawed and resuspended in a mixture of 92% NKM and 5.5% 2× DMEM (Dulbecco’s Modified Eagle Medium) to generate the cell master mix. NKM consisted of KnockOut^TM^ DMEM (KO-DMEM, Gibco, ThermoFisher Scientific, USA) supplemented with 10% fetal bovine serum (FBS, Sigma-Aldrich, USA) and 1% GlutaMAX (Gibco, ThermoFisher Scientific, USA), whereas 2× DMEM was prepared from a sterile-filtered 10× DMEM (Sigma-Aldrich, USA) stock combined with 50% FBS and 5% penicillin–streptomycin.

The custom EHT preparation assembly (BioGenium Microsystems Oy, Tampere, Finland) consisted of a 1-well Sample Cup, a removable Gel-Mould, and an EHT-LID containing one pair of flexible pillars (Diameter 0.18 mm, length 5 mm, center-to-center distance between the pillars 5 mm). Sterile 2% UltraPure^TM^ Agarose (Sigma-Aldrich, USA) was used to prepare EHT mold in advance. During agarose casting, the Gel-Mould formed a mold cavity at the center of the 1-well Sample Cup. After the agarose had solidified completely, the Gel-Mould was removed, and the EHT-LID was positioned so that the pillar pair extended into the agarose mold cavity. Immediately before casting, fibrinogen from bovine plasma (Sigma-Aldrich, USA) and thrombin from bovine plasma (Sigma-Aldrich, USA) were added to the cell master mix to achieve final concentrations of 2 mg/ml fibrinogen and 0.6 U/ml thrombin. The resulting cell-hydrogel mixture was then pipetted carefully into the agarose mold cavity around the pillar pair (Fig. 1a). The EHT construct was incubated for 20 min at 37 °C to allow complete fibrin polymerization. Following polymerization, the EHT-LID carrying the EHT construct was lifted from the agarose mold and transferred to a separate 1-well Sample Cup containing EHT medium. The EHT medium consisted of KO-DMEM supplemented with 10% FBS, 10 μg/ml insulin solution human (Sigma-Aldrich, USA), 33 μg/ml aprotinin from bovine lung (Sigma-Aldrich, USA), and 1% penicillin–streptomycin. Tissues were maintained for two weeks, with the medium refreshed three times per week.

**Fig. 1.**
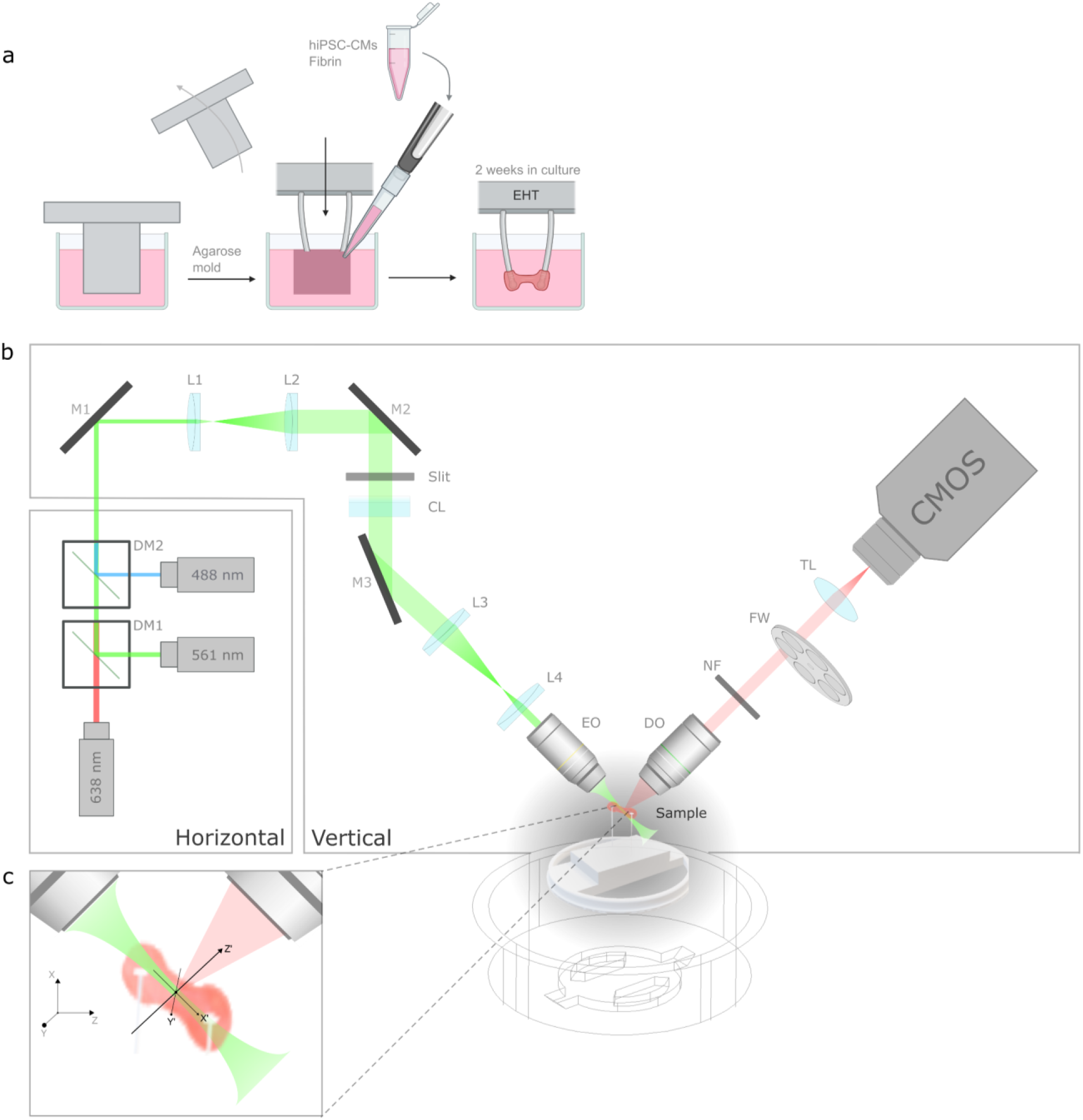
Schematic illustration of iSPIM platform for EHT imaging. **a** Illustration of molding and casting the engineered heart tissue (EHT) construct. The EHT is formed with human-induced pluripotent stem cell-derived cardiomyocytes (hiPSC-CMs) in fibrin hydrogel and cultured for 2 weeks before imaging. **b** Optical schematic of the inverted selective plane illumination microscopy (iSPIM) system with the horizontal and vertical parts indicated separately, combined with the water-immersion sample holder. **c** Scheme of the reference systems; the XYZ of the microscope motor controls and the X’Y’Z’ of the illuminated and detected section of the sample. DM1-2: dichroic mirror, M1-M3: mirror, L1-L4: lenses, CL: cylindrical lens, EO: excitation objective, DO: detection objective, NF: notch filter, FW: filter wheel, TL: tube lens

### 2.4 Inverted selective plane illumination microscope imaging platform

An iSPIM platform was developed for volumetric imaging of EHTs. The schematics of the iSPIM platform together with EHT sample holder are presented in Fig 1. The inverted configuration supported optimal positioning of the two-pole EHTs with the long axis of the tissue in between the microscope objectives (Fig 1b). The iSPIM system consists of an excitation arm and an orthogonally positioned detection arm. The EHT is mounted on a sample holder attached to motorized linear (Precision Linear Stage M-404.1PD, Physik Instrumente, Germany) and rotation (Precision Rotation Stage M-060.PD, Physik Instrumente, Germany) stages for precise and volumetric image acquisition. Manual positioning along X-axis was conducted adjusting the vertical position of the sample (L490/M, Thorlabs Inc, USA) (Fig 1c). During imaging, the EHT was immersed inside a water bath alongside both objectives.

In the illumination path, three laser lines, 488, 561, and 638 nm (Cobolt 06-series, Hüber Photonics) were combined using two dichroic mirrors, DM1 (Di03-R561-t1, Semrock, USA) and DM2 (Di03-R488-t1, Semrock, USA). The 638 nm laser passed through both DM1 and DM2, the 561 nm laser was reflected by DM1 and transmitted through DM2, and the 488 nm laser was reflected by DM2. The combined beam was then directed by a mirror (M1, BB1-E02, Thorlabs Inc, USA) and expanded using a Kepler telescope consisting of lenses L1 (f=30 mm, AC254-030-A-ML, Thorlabs Inc, USA) and L2 (f=60 mm, AC254-060-A-ML, Thorlabs Inc, USA). The expanded beam was subsequently reflected by a second identical mirror (M2) and passed through a rectangular slit (VA100CP/M, Thorlabs Inc, USA). A cylindrical lens CL (f=50 mm, ACY254-050-A, Thorlabs Inc, USA) then focused the beam in one dimension to generate the light-sheet. Finally, the shaped beam was reflected by a third mirror (M3) and relayed through a second telescope consisting of lenses L3 (f=75 mm, AC254-075-A-ML, Thorlabs Inc, USA) and L4 (f=30 mm, AC254-030-A-ML, Thorlabs Inc, USA) to generate a thinner light-sheet at the same plane. The light-sheet was then focused onto the EHT using a 10X/0.3 numerical aperture (NA) water-immersion excitation objective (EO) (UMPLFLN 10XW, Olympus, Japan).

The emitted fluorescence from the sample was collected using a 20X/0.5 NA water-immersion detection objective (DO) (UMPLFLN 20XW, Olympus, Japan). The emitted signal was transmitted through a notch filter (NF, StopLine quad-notch filter 405/488/561/635 nm, Semrock, USA), while the excitation wavelength was selectively rejected. The filtered signal was directed to a motorized filter wheel (FW103 Series, Thorlabs Inc, USA) equipped with wavelength-specific emission filters (488 nm: BLP01-488R-25, Semrock, USA; 561 nm: FBH590-10, Thorlabs Inc, USA; 638 nm: BLP01-633R-25, Semrock, USA) to further filter out any remaining excitation laser and non-specific fluorescence before detection. Finally, the fluorescence signal was relayed through a tube lens (TTL165-A, f=165 mm, Thorlabs Inc, USA) and imaged onto a digital scientific complementary metal-oxide-semiconductor (sCMOS) camera (ORCA-Flash4.0 V3, Hamamatsu, Japan). The image size of 2048 x 2048 pixels with effective pixel size of 325 nm x 325 nm was acquired. The 3D image stacks were acquired by translating the sample toward the excitation objective using a motorized linear stage, while sequential optical sections were recorded by the camera (Fig 1c). The iSPIM system was controlled using an Inscoper Imaging Solution controller and software (Inscoper SAS, France).

### 2.5 Immunocytochemical staining and iterative indirect immunofluorescence

For fluorescence imaging, EHTs were labeled for cTnT, Cx43, and nuclei to visualize sarcomere structures, intercellular gap-junctions, and cell nuclei, respectively. All fixation and staining procedures were performed as previously described [45] while the EHTs remained attached to the poles. Briefly, EHTs were fixed with 4% paraformaldehyde (PFA, Acros Organics, Belgium) in phosphate buffered saline (PBS, EuroClone, Italy) for 1 h at room temperature. Following two 10 min washes with PBS, the EHTs were blocked overnight at +4 °C with the blocking solution containing 10% normal donkey serum (NDS, Millipore, USA), 1% bovine serum albumin (BSA, Sigma-Aldrich, USA), and 0.1% TritonX-100 (Sigma-Aldrich, USA).

After a 5 min wash in 1% NDS, 0.1% Triton X-100, and 1% BSA in PBS, the tissues were incubated with primary antibodies for three days at +4 °C on a shaker. The primary antibody solution contained mouse anti-connexin 43 in 1:300 (Millipore, CAT# MAB3068, RRID:AB_2110187) and rabbit cardiac troponin T in 1:150 (Abcam, Cat# ab92546, RRID:AB_10585593) diluted in 1% NDS, 0.1% Triton X-100, and 1% BSA in PBS. After primary antibody incubation, the EHTs were washed with 1% BSA in PBS sequentially for 10 min, 7 h, overnight, 7 h, and a final overnight wash.

Secondary antibody incubation was performed overnight at +4 °C in 1% BSA in PBS with Alexa Fluor^TM^ 488 anti-Mouse (Invitrogen, CAT# A21202, 1:800 dilution) and Alexa Fluor^TM^ 568 anti-Rabbit (Invitrogen, CAT# A10042, 1:800 dilution). Following secondary antibody incubation, the EHTs were washed with PBS for three sequential washes for 10 min, 1 h, and 10 min. Nuclei were stained using NucSpot (NucSpot Live 650, Biotium, CAT# 41034, in 1:1000) during the 1 h wash with PBS.

In addition to the regular immunostaining protocol, IT-IF was applied as described in [46] using cTnT staining. Briefly, two consecutive rounds of immunostaining were applied to generate a 2X iterative staining sample, with the blocking step omitted during the second staining cycle. Parallel EHT samples stained with a conventional single-round immunostaining protocol were used as controls and are hereafter referred to as 1X tissue.

### 2.6 Experimental point spread function measurement

For characterization of the iSPIM system, 500 nm fluorescence beads (FluoSpheres™, Invitrogen, USA) were embedded in 1% agarose gel (Sigma, USA) and inserted into an optically transparent fluorinated ethylene propylene (FEP, Adtech Polymer Engineering) tube. The bead sample was excited using 561 nm laser line and imaged through corresponding emission filter. Images were acquired with an effective pixel size of 325 nm x 325 nm in XY plane, Z step size of 200 nm, and an exposure time of 100 ms. Laser power was adjusted to maximize the fluorescence signal without saturation.

Experimental point spread function (PSF) was measured from the fluorescent bead images to evaluate the resolution of the iSPIM system. Huygens Essential software version 25.10 (Scientific Volume Imaging, The Netherlands) PSF Distiller was used for calculating the experimental PSF. The theoretical PSF was calculated with the same software Gaussian light-sheet mode with the following parameters: excitation wavelength 561 nm, detection objective NA of 0.5, light-sheet thickness of 10 µm, and excitation direction from left of the sample. The experimental and theoretical PSFs were compared in MATLAB R2024b (The MathWorks Inc., USA). The full width at half maximum (FWHM) was used to compare experimental and theoretical PSFs.

### 2.7 Zebrafish imaging for iSPIM workflow validation

The zebrafish line Tg(fli1:EGFP)y1 was used to validate the imaging and image processing workflow of the iSPIM system. The transgenic line expresses green fluorescent protein (GFP) in the endothelial cells throughout the vasculature under control of the *fli1* promoter [47]. At 1 dpf, embryos were dechorionated, screened and transferred to 0.0045% PTU (1-phenyl-2-thiourea) -containing water. At 3 dpf, embryos were euthanized (overdose of MS-222) and fixed in 10% neutral buffered formalin prior to imaging. Fixed embryos were embedded in 1% agarose gel into a FEP tube and imaged using 488 nm laser line. 3D stack images were acquired with a Z step size of 500 nm.

### 2.8 Volumetric imaging of engineered heart tissues

The iSPIM platform was used for volumetric imaging of EHTs to assess CM alignment, gap-junction distribution, and cellular content. The fluorescence signals of Cx43, cTnT, and nuclei were imaged using the corresponding excitation/emission channels with 100 ms exposure time, adjusting the laser power to avoid photobleaching and saturation. The images were acquired with 2048 x 2048 pixels, with a voxel size of 325 nm x 325 nm x 1 µm. For IT-IF studies, imaging was performed starting with the 2X iterated tissue, maintaining the same imaging parameters for single-round immunostaining protocol control. Tissue sections in between the poles were focused on and the full tissue thickness (approx. 300 µm) was imaged in less than 60 s per channel. Fiji [48] was used to visualize maximum intensity projections (MIP), whereas Napari viewer was used for volumetric visualization of the EHT. For full EHT visualization, six adjacent imaging positions with a 10% overlap were aligned and stitched using the Grid/Collection Stitching plugin in Fiji [49]. Supplementary Video S1 was generated in Imaris (v11.0.1, Oxford Instruments, Switzerland) from the same tissue volume after fivefold downsampling in all dimensions (X, Y and Z). Furthermore, for more detailed volumetric visualization, two adjacent imaging positions with a 10% overlap were similarly aligned and stitched and viewed with volume rendering using a 5% attenuated MIP approach to give an overview in 3D of the EHT. For comparison, one EHT was processed and imaged with iSPIM as described, and then detached, cleared and imaged using confocal microscopy (Additional file 1: Supplementary Materials and Methods).

### 2.9 3D image processing, deconvolution, and analysis

To correct for the oblique acquisition geometry inherent to inverted light-sheet microscopy, raw iSPIM image stacks were deskewed prior to further image processing and analysis. Deskewing was performed using an affine transformation [50], with a light-sheet angle of 50° and the acquisition Z step size used as input parameters. The resulting deskewed image stacks were subsequently deconvolved using the light-sheet module in Huygens Essential software to improve image contrast and resolution.

For deconvolution using theoretical PSF, imaging parameters were specified in the software to generate the theoretical PSF (voxel size: 325 nm x 325 nm x 766 nm; excitation/emission wavelengths for Cx43, cTnT, and nuclei in nm: 488/520, 561/590, 638/650, respectively; NA of the DO: 0.5; refractive index of the sample media: 1.338 with no refractive index mismatch; Gaussian light-sheet thickness:10 µm; imaging direction: left). For deconvolution using experimental PSF, the measured PSF described in section 2.6 was used. In both cases, the same deconvolution parameters were applied, including Classic deconvolution algorithm, signal to noise (SNR) ratio of 30, number of iterations of 75, and quality threshold of 0.01. The light-sheet module deconvolution considers spatial variation of the PSF true to Gaussian light-sheet.

To compare the effect of deconvolution using theoretical and experimental PSFs, zebrafish embryo image stacks acquired as described in section 2.7 were processed using both approaches and compared with the corresponding raw datasets. To assess changes in image resolution and contrast, intensity line profiles were extracted from selected vascular structures in both lateral and axial direction using the Plot Profile function in Fiji. The effect of deconvolution approaches to image contrast was determined from non-resolvable structures of the raw dataset by calculating the ratio between signal peak maximum through minimum. The effect of deconvolution approaches to resolution was assessed by calculating the FWHM from a fitted Gaussian curve to the intensity line profile.

### 2.10 Analysis of indirect iterative immunofluorescence

The change in fluorescence signal intensity and signal-to-background ratio (SBR) was analyzed during the IT-IF process to assess the contribution of specific cTnT antibody binding relative to non-specific background signal. The average fluorescence intensity was measured from a 200 x 200-pixel region of interest (ROI) located at the center of the EHT throughout tissue depth. SBR was determined using Labelme toolkit [51] for manually labeling in-focus CMs throughout the Z-stack. Measurements were performed on every 25th image slice, corresponding approximately to 19 µm intervals along the imaging depth. The labeled cell regions were converted into a binary mask. Each mask was dilated by 15 pixels and the original mask was subtracted from dilated mask to generate a surrounding ring-shaped background region for SBR measurements. The same SBR analysis was conducted on the deconvolved images of the IT-IF stained EHTs. All image processing and quantitative analysis were performed using custom Python scripts.

### 2.11 3D segmentation of nuclei and registration for cardiomyocytes

For quantification of tissue composition, nuclei were used as seeds to define individual cells within the EHT. 3D nuclei segmentation was performed using ilastik (version 1.4.1) [52]. First, the Pixel Classification workflow was used to assign voxels into background and foreground classes. Two foreground classes were defined to distinguish the slightly brighter nuclei edges from the nuclear interior. The classifier was trained using manually annotated representative slices in XY planes together with sparse annotations in YZ and XZ planes. The trained model was subsequently applied to 3D nuclei datasets, and the resulting voxel probability maps were exported in HDF5 format.

Next, the Object Classification workflow was used to segment individual nuclei in 3D. The voxel probability maps and the original images were used as inputs. Hysteresis thresholding was used with a core threshold of 0.8, a final threshold of 0.6, and a smoothing factor of 3 in all directions. Objects smaller than 300 voxels were removed, and sparse manual labeling was added for the classifier. The trained object classifier was then applied to all datasets in batch mode. Segmented nuclei were exported originally as HDF5 files and converted to a multipage TIFF file using the Data Conversion workflow.

Segmented nuclei were classified as cardiomyocyte-positive (CM+) or cardiomyocyte-negative (CM-) based on the cTnT channel. Intensity threshold was determined by Otsu thresholding, and the average cTnT intensity associated with each segmented nucleus was calculated and used for classification. Napari was used to visualize the spatial localization of the CM+ nuclei within a representative EHT volume. A total of 11 images from n=5 different EHTs were segmented and classified, and the % of CM+ nuclei of total segmented nuclei was calculated.

To evaluate the depth-dependent proportion of CM+ nuclei, the percentage of CM+ nuclei was calculated from tissue surface to tissue core using 20 µm thick depth intervals. The central tissue slice was defined in Fiji, and the centroid coordinates of segmented nuclei were used for depth-dependent registration in 3D. Furthermore, tissue heterogeneity was evaluated by dividing the representative EHT stack into six subsections along the Y-axis (approximately 110 µm long) and comparing the depth-dependent percentages of CM+ nuclei between subsections.

### 2.12 Cardiomyocyte orientation analysis

CM alignment throughout the EHT was quantified from cTnT fluorescence image stacks using texture analysis calculated from intensity gradient orientations. Regions corresponding to tissue boundaries were excluded from the orientation analysis to minimize edge effects. The orientation analysis was performed on full 3D image stacks in a slice-by-slice manner.

Structure tensor components for each image slice were computed using the structure_tensor function from the scikit-image Python library (version 0.24.0) [53]. The image was first smoothed with a Gaussian filter (σ=3), and a Gaussian kernel size σ=15 was selected for computing the structure tensor. The local orientation, coherence and energy were calculated from eigenvalues extracted from the structure tensor, similarly to the OrientationJ plugin for Fiji (Biomedical Imaging Group, EPFL) [54]. Orientations with coherence value below 0.2 were excluded from further analysis to reduce the effect of random noise. To account for the inherent 180° symmetry of orientations, all orientation angles were restricted to the range [0, 180].

Orientation histograms generated from the 3D image volumes were normalized to probability densities by dividing by the number of pixels used and histogram bin width, thereby eliminating the influence of differences in cTnT pixel counts between image slices. Orientation maps were visualized using OrientationJ plugin in Fiji, where hue color represents the local orientation angle.

Three separate sections of the same EHT were quantified for mapping tissue heterogeneity. Each image slice from each tissue section was summarized by calculating the dominant direction using the axial circular mean of orientations, with weights given by the product of coherence and energy values. The dominant directions were compared between the tissue sections.

### 2.13 Statistical analysis

For all statistical tests, data normality was assessed using the Kolmogorov-Smirnov test and visual inspection of Q-Q plots. Non-parametric Mann-Whitney U test was used to test statistical differences between groups. Bonferroni correction was applied to account for multiple comparisons. The results were presented as median (25th–75th percentile). P-values <0.05 were considered statistically significant. All statistical analysis were performed using IBM SPSS Statistics version 30 (SPSS, USA). In IT-IF analysis, all data were normalized to the 1X control tissue from the same experiment prior to statistical testing.

## 3. Results

### 3.1 Characterization and validation of the iSPIM imaging platform

The imaging performance of the developed iSPIM platform was assessed by characterizing the PSF using sub-resolution fluorescent beads. The orthogonal slice views of the beads across partial field of view (FOV) and of an average single bead used as the experimental PSF are shown in Fig 2a and b, respectively, providing an overview of bead distribution within the imaged volume. The beads appear circular in lateral planes, while elongated along the axial direction (see Fig 2b and c). The experimental PSF profiles along all axes were obtained from multiple fluorescence beads and are presented as an average PSF together with the theoretical PSF in Fig 2d. The measured PSF was broader than the theoretical PSF in all three dimensions, with lateral FWHM approximately twice the theoretical values (1.1 µm vs. 0.59 µm) and an axial FWHM approximately twice the theoretical values (9.6 µm vs. 3.96 µm). These measurements demonstrate anisotropic spatial resolution of the iSPIM platform, with higher resolving power in the lateral plane than along the axial direction.

**Fig. 2.**
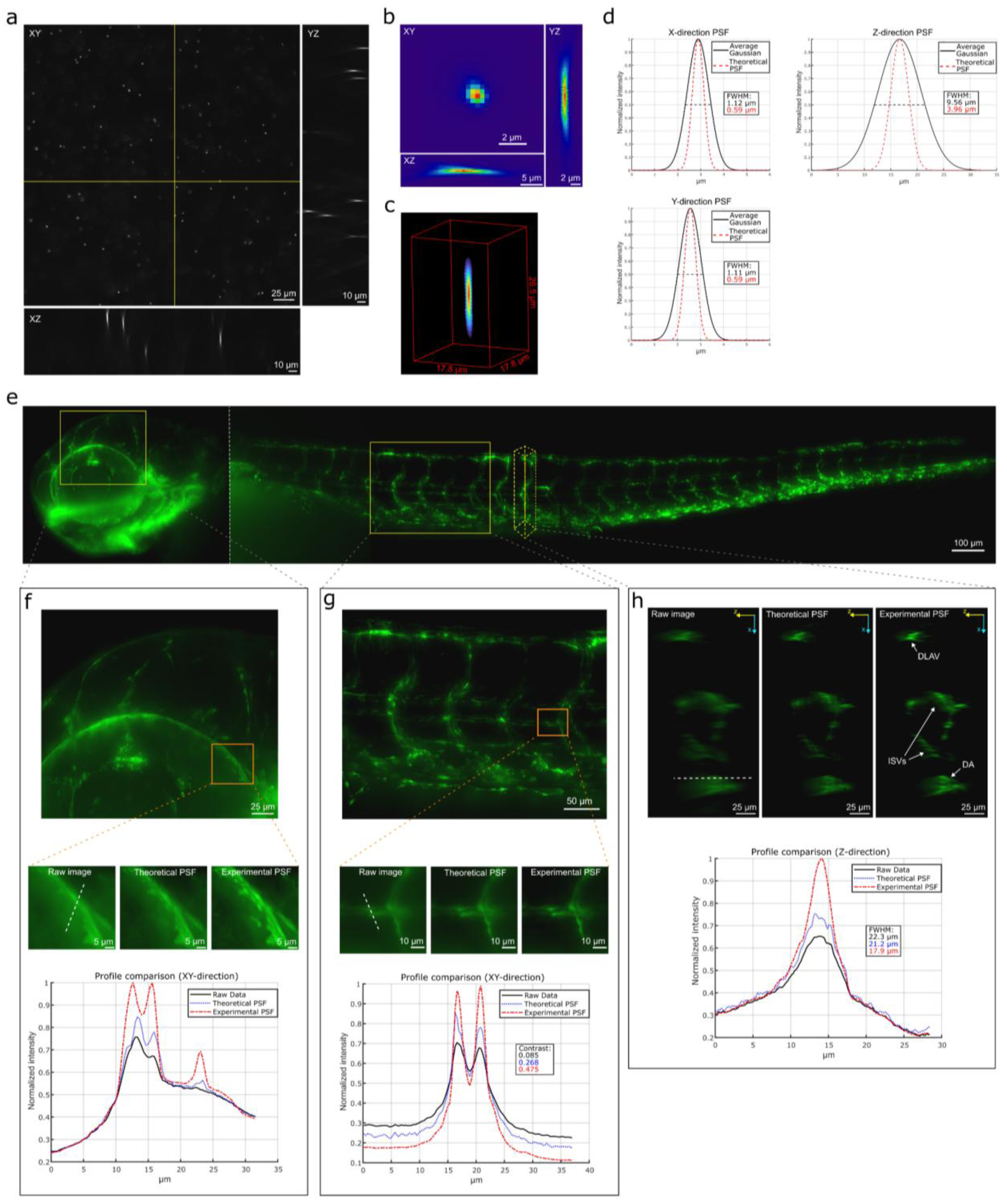
iSPIM characterization for structural imaging. **a** Orthogonal views of a larger field of view (FOV) of 500 nm fluorescent beads. **b** Experimental point spread function (PSF) orthogonal views. **c** Volume rendered view of the PSF. **d** Fitted Gaussian profiles both to the experimentally measured PSF and theoretically derived PSF plotted for all axes X, Y and Z with measured full width half maximum (FWHM) marked with corresponding colors. **e** Zebrafish embryo maximum intensity projections (MIP) of the tail and head area. **f** Eye vascular structures of the embryo, with enlarged regions and corresponding line profiles demonstrating that experimental PSF deconvolution resolves vascular features more clearly than both the raw image and theoretical PSF deconvolution. **g** Tail vascular structures with enlarged regions and corresponding line profiles showing improved image contrast after deconvolution, as quantified in the inset, compared to the raw image. **h** Axial XZ MIP views of a tail section with line profiles and FWHM measurements demonstrating improved axial resolution using experimental PSF deconvolution. DLAV: dorsal longitudinal anastomotic vessel, ISVs: intersegmental vessels, DA: dorsal aorta

The well-characterized vasculature of a 3 dpf zebrafish embryo was imaged to further validate the imaging capabilities of the iSPIM system. GFP-tagged vascular endothelial cells were clearly visualized throughout the embryo (Fig. 2e). Subsequent deconvolution of the 3D image dataset using both theoretical and experimental PSFs further improved the visualization of selected vascular structures (Fig 2f and g). Both methods enhanced image contrast and resolution around the eye (Fig 2f), with the experimental PSF producing the strongest improvement. The improvement in image quality was further confirmed by the contrast measurement between two closely spaced vascular structures in the trunk (Fig 2g). Experimental PSF deconvolution produced the highest contrast (0.475), followed by theoretical PSF deconvolution (0.268), whereas the raw image exhibited substantially lower contrast (0.085). Similar improvements were observed in axial view of the zebrafish vasculature in the trunk (Fig 2h). Deconvolution enhanced identification of dorsal longitudinal anastomotic vessel (DLAV), intersegmental vessels (ISVs), and dorsal aorta (DA). In addition, apparent thickness of DA wall in axial direction decreased from 22.3 µm in the raw image to 21.2 µm and 17.9 µm following theoretical and experimental PSF deconvolution, respectively, indicating reduced axial blurring and improved structural definition. Together, these results demonstrate that the iSPIM platform, combined with experimental PSF deconvolution, enables high-quality volumetric structural imaging. Consequently, experimental PSF deconvolution was selected for all subsequent image processing and structural analyses.

### 3.2 Signal enhancement in EHTs using iterative indirect immunofluorescence

To improve the fluorescence signal for iSPIM imaging of intact EHTs, IT-IF was applied. An additional round of cTnT labeling was performed to generate the 2X stained tissue, and both 1X and 2X tissues were imaged using identical acquisition settings as illustrated in the workflow shown in Fig 3a. The mean intensity and SBR measurements used for evaluating IT-IF performance are illustrated in Fig 3b. IT-IF resulted in a significant increase in cTnT fluorescence intensity in 2X tissue compared to 1X conventional staining (5.17 [3.77-6.40] vs. 1.02 [0.77-1.22], respectively), with the 2X iterated EHTs exhibiting more than a fivefold increase in intensity (Fig 3c; p<0.001). An increase in SBR was also observed in the 2X iterated EHTs compared with the 1X tissues (1.04 [0.96-1.31] vs. 0.99 [0.89-1.09], respectively) which was statistically significant (Fig 3d; p<0.001). Processing the images with deconvolution, this 5% increase in the median SBR was further increased to 7.5% (1.04 [0.90-1.35] vs. 0.97 [0.84-1.09], respectively), remaining as a significant increase (Fig 3d; p<0.001). These results indicate that IT-IF increased not only the overall fluorescence intensity but also the specificity of cTnT signal relative to the background. The SBR remained relatively stable through tissue depth, although increased variation was observed in deeper tissue regions (Fig 3e). Together, these results demonstrate that IT-IF substantially enhances fluorescence signal and SBR in intact EHTs, enabling improved structural imaging with iSPIM throughout the tissue volume.

**Fig. 3.**
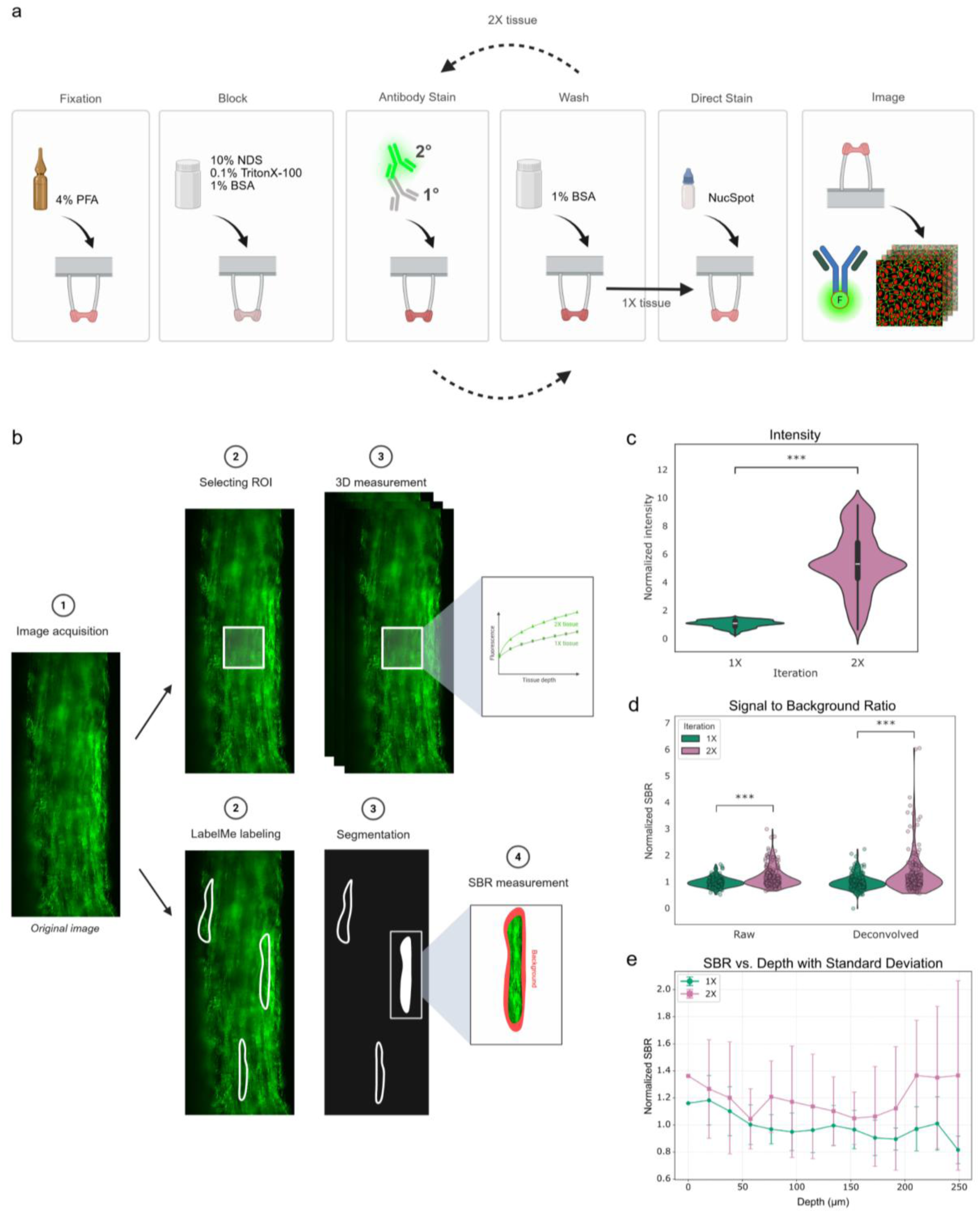
Improving fluorescence signal of EHTs with IT-IF. **a** The overall workflow of iterative indirect immunofluorescence (IT-IF) presented as a schematic. **b** The methods of analysis are presented for comparison between conventional immunostaining protocol (1X) and with two-round iteration staining (2X) in raw intensity (upper row) and in signal to background ratio (SBR) (lower row). **c** IT-IF significantly increased the intensity level of 2X engineered heart tissues (EHTs) for cardiac troponin T (cTnT) labeling. **d** The SBR was increased significantly in both raw and deconvolved images of cTnT labeling, indicating labeling specificity to the target. **e** The SBR of deconvolved images plotted through tissue depth, measuring at every 25th slice in the EHT volume, starting from the tissue surface. Standard deviation is plotted as positive and negative error bars. Statistical significance: ***p<0.001. n=3 independent experiments, intensity comparison n=8 sections and n=10 sections for 1X and 2X area measurements, respectively, and SBR comparison n=203 and n=216 cells selected for 1X and 2X, respectively

### 3.3 3D visualization of engineered heart tissue

An overview of the EHT is presented to illustrate the overall tissue architecture and provide context for subsequent structural analyses. Imaging was acquired from the tissue region between the poles, with a full tissue visualization for cTnT and nuclei (Fig. 4a) and in more detail in supplementary video (Video S1). Individual slices from top, middle, and bottom of a 110 µm thick EHT volume allowed visualization of Cx43, cTnT, and nuclei clearly throughout the tissue (Fig 4b). CMs, identified by cTnT labeling, appeared visually aligned with the major axis of the tissue in all depths. Cx43 appeared as dots, marking gap-junctions throughout the tissue. Visually high density of nuclei was also visible throughout the imaged EHT volume. A MIP of the full tissue section showed an overview of the tissue organization (Fig 4c). Furthermore, the 3D rendering generated from two stitched image volumes, acquired from the center area of the EHT, revealed continuity of the aligned CM structures across a 1.28 mm long tissue volume together with the spatial distribution of Cx43 and nuclei (Fig 4d). These results demonstrate that iSPIM enables volumetric imaging of EHTs, providing visualization of cardiac structures from subcellular to tissue scale.

**Fig. 4.**
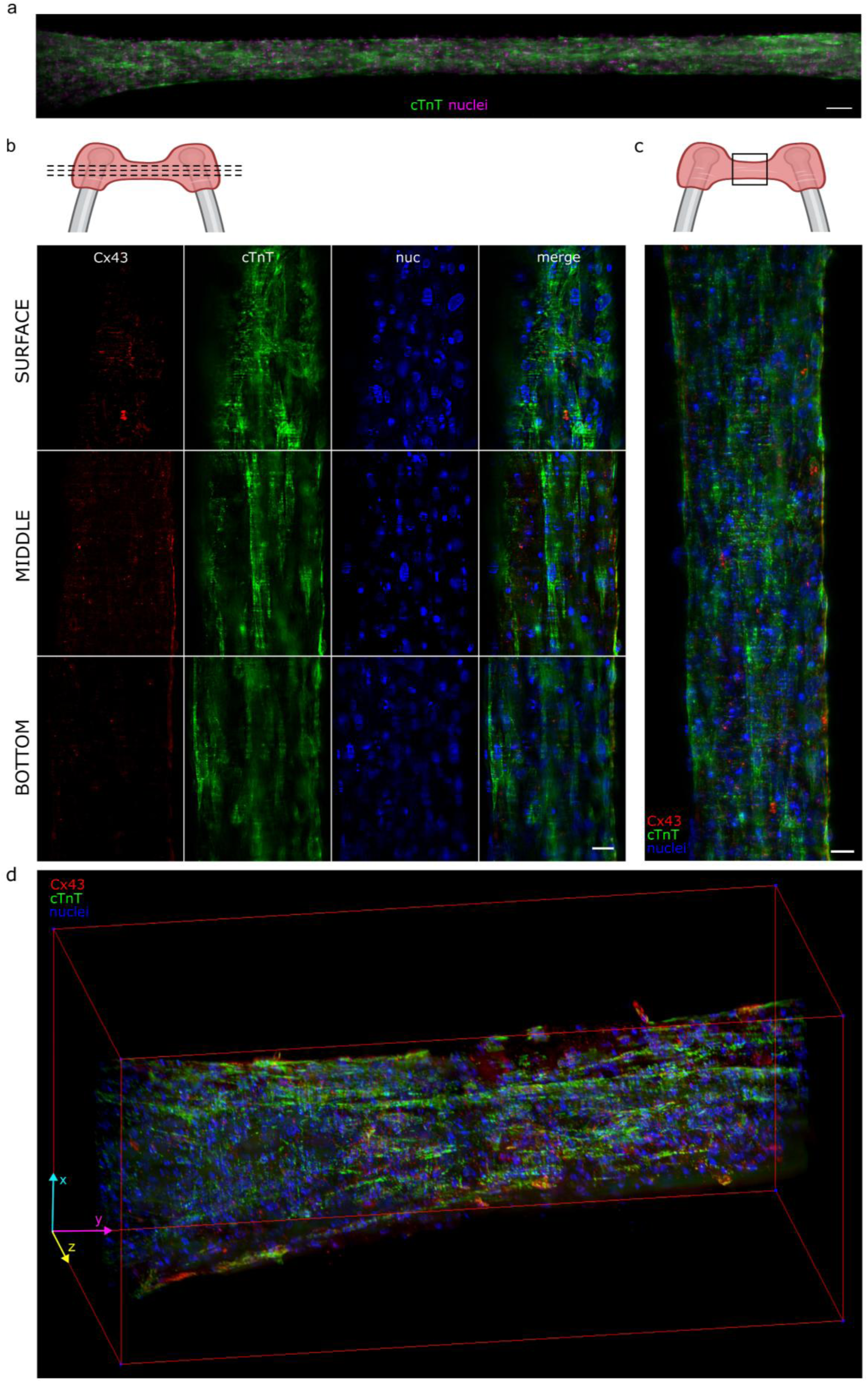
Volumetric imaging of EHTs with iSPIM platform. **a** Full EHT tissue visualized with maximum intensity projection (MIP) between the poles. The cardiac-specific troponin T (cTnT) is shown in green and the nuclei in magenta (scale bar: 100 µm). **b** 2D slices from an EHT volume visualized for cTnT in green, connexin 43 (Cx43) in red, and nuclei in blue at top, middle, and bottom depth of the imaged volume (scale bar: 25 µm). **c** A MIP of the full tissue volume, labeled with cTnT, Cx43, and nuclei (scale bar: 25 µm). **d** Volume rendering view of the 3D EHT section between the poles.

### 3.4 Cardiomyocyte content and distribution in EHTs

CM content within the EHT was quantified by classifying segmented nuclei as CM-positive (CM+) or CM-negative (CM-) based on cTnT labeling. Nuclei were successfully segmented throughout the imaged EHT volume (Fig 5a) and registered to the cTnT channel using intensity-based thresholding for CM classification (Fig 5b). Using this approach, 71.4% of nuclei were classified as CM+ on average across five replicate tissues (Fig 5c). In addition, the distribution of CM+/CM-nuclei was assessed from the surface of an EHT to the tissue core, rising from approximately 40% near the surface to around 60% at the core (Fig 5d). To assess regional differences in CM content, the segmented tissue volume was divided into six subsections (Fig 5e). Differences in the distribution of CM+ nuclei were apparent between subsections and were further quantified by analysis of CM+/CM-content across the EHT depth (Fig 5f). These findings demonstrate that volumetric imaging combined with 3D nuclei segmentation and registration enables quantitative assessment of CM content and spatial heterogeneity across the EHT volume.

**Fig. 5.**
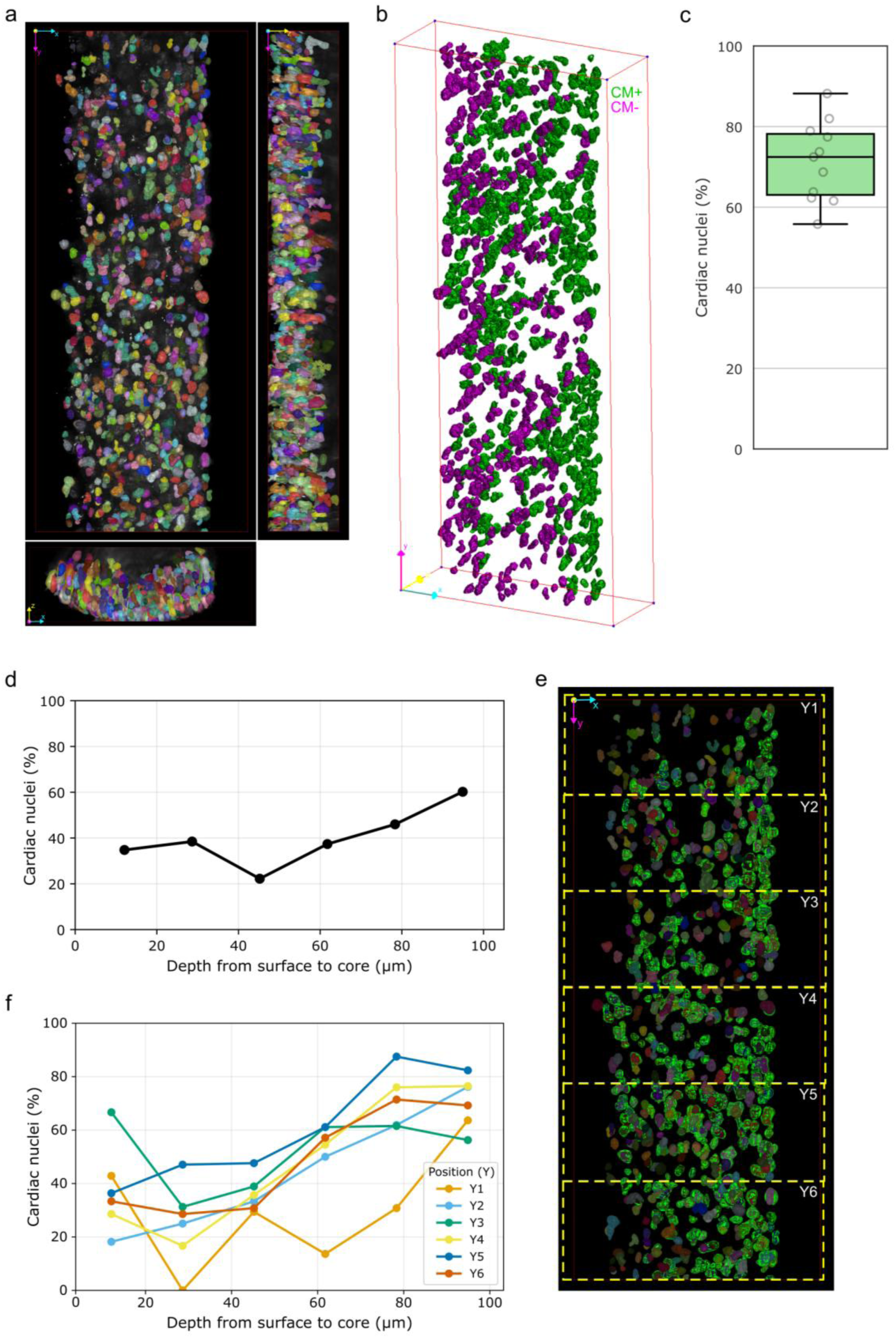
3D segmentation and registration of cardiac nuclei. **a** 3D segmentation of nuclei within an engineered heart tissue (EHT) visualized in XY, YZ and XZ views (isotropic scaling in volume rendering). **b** Intensity-based registration of the segmented nuclei to cardiomyocyte (CM+, visualized in green) or not cardiomyocytes (CM-, visualized in magenta), viewed with volume rendering. **c** CM+ nuclei percentage quantified from 11 imaged tissue sections (n=5 separate tissues). **d** Overall percentage of CM+ nuclei through the tissue depth from tissue surface (0 µm) to core (100 µm), binned with 20 µm sections. **e** Front view of the segmented and registered nuclei (CM+ shown in green), where the six tissue subsections (Y1-Y6) are indicated. **f** The six subsections corresponding to e plotted for CM+ nuclei percentage through the tissue depth

### 3.5 3D alignment of cardiomyocytes in EHTs

CM alignment was quantified throughout the tissue volume from cTnT labeled images to characterize the 3D organization of the EHT. CM alignment was evaluated on a slice-by-slice basis to preserve depth-dependent differences throughout the tissue volume (Fig 6a). The resulting orientation distributions exhibited a pronounced peak around 90° throughout most of the tissue depth, indicating a high degree of alignment along the longest axis of the tissue (Fig 6b). The distributions became broader in the deeper parts of the tissue. To assess whether alignment varied between different regions of the tissue, dominant orientation was evaluated for each slice in three consecutive EHT. Significant differences in alignment were detected between the sections (Fig 6c; p<0.001), demonstrating spatial variation in CM alignment across the EHT (section 1: 89.7° [88.4-90.4] and section 2: 89.6° [86.7-91.1] vs. section 3: 87.6° [83.7-89.1]). These findings highlight the importance of depth-resolved analysis for accurate characterization of CM organization and regional heterogeneity in 3D EHTs.

**Fig. 6.**
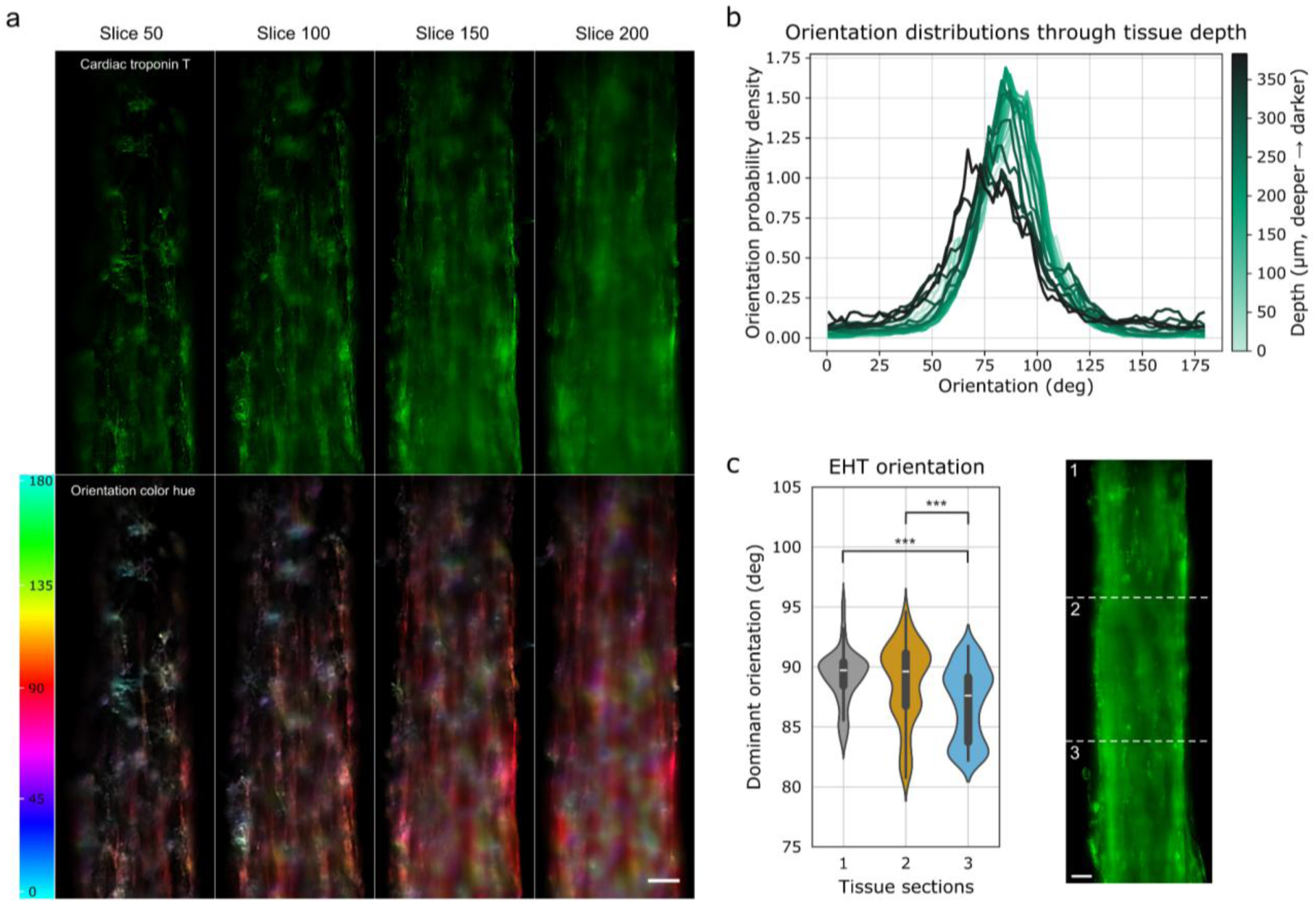
Characterization of CM alignment through depth in EHT. **a** Representative image slices of cardiac troponin T (cTnT) channel and the orientation analysis visualized with a color hue (scale bar: 50 µm). **b** Normalized probability density of cardiomyocyte (CM) alignment orientations through the tissue depth, with darker color on the plot indicating greater imaging depth in the tissue. **c** Depth-resolved orientation distributions from three sections of the same EHT tissue with significant difference detected between the third section to the other two sections. The tissue sections are visualized as maximum intensity projection (MIP) view with dashed white lines and numbers corresponding to each section (scale bar: 50 µm). Statistical significance: ***p<0.001. n=510 slices through the tissue for each section

For comparison, the same EHT was imaged first with iSPIM and subsequently detached, optically cleared, and imaged with confocal microscopy. Although the same tissue regions appeared visually similar with both methods, dominant-angle orientation analysis showed substantial differences between the iSPIM and confocal images (see Additional file 1: Supplementary Results and Figure S). These findings demonstrate the feasibility of assessing EHT organization directly in intact construct using iSPIM.

## 4. Discussion

In this study, we demonstrate that the developed iSPIM platform, combined with IT-IF staining and an image analysis pipeline, enables volumetric imaging and 3D cellular and subcellular structural characterization of EHTs. The developed workflow enables quantitative assessment of cardiomyocyte alignment, cellular composition, and spatial heterogeneity throughout the tissue volume. To our knowledge, such quantitative characterization of intact EHTs has not previously been reported. These capabilities extend beyond EHTs, providing a broadly applicable approach for in situ, quantitative 3D characterization of engineered tissues with sufficient optical transparency for volumetric imaging.

Volumetric imaging of intact EHTs remains challenging due to tissue thickness and the geometry of the construct. The presented iSPIM configuration enabled imaging of intact EHTs between the excitation and detection objectives, facilitating optical sectioning while preserving native tissue architecture. The imaging performance of the system was assessed through PSF characterization. The measured lateral and axial resolutions were 1.1 µm and 9.6 µm, respectively, comparable to previously reported light-sheet microscopy systems with similar optical configurations [31,55]. As expected for single-view SPIM, the system exhibited anisotropic resolution, with higher resolving power in the lateral plane than along the axial direction. This is characteristic of Gaussian light-sheet microscopy, where the axial resolution is influenced by both light-sheet thickness and detection optics [29,56]. Consequently, structures appeared elongated along the optical axis. Experimental PSF-based deconvolution reduced this effect and improved image contrast and structural detail. Thus, the achieved image quality was sufficient for reliable volumetric visualization and quantitative structural analysis of intact EHTs.

Achieving sufficient fluorescence signal and contrast throughout the tissue volume is critical for volumetric imaging and quantitative analysis. In the present study, IT-IF substantially enhanced cTnT labeling, resulting in a more than fivefold increase in fluorescence intensity together with a significant improvement in SBR. The increase in SBR was particularly important, as it suggests that the enhanced fluorescence signal primarily originated from specific cTnT labeling rather than increased non-specific binding, consistent with previous findings [46]. Antibody penetration nevertheless remains a known challenge in large 3D tissue constructs and may contribute to labeling variability in deeper tissue regions [57]. Consistent with this, a modest decrease in SBR with imaging depth was observed in both 1X and 2X tissues, reflecting the increasing contribution of out-of-focus signal to the measured background. While tissue clearing may further improve imaging depth and contrast by reducing light scattering [58–60], it requires additional sample preparation and may alter tissue morphology [61]. Furthermore, cardiac tissue clearing remains less established than corresponding approaches in neuroscience [60]. However, further optimization of these approaches may improve depth-resolved structural assessment of EHTs. Nevertheless, sufficient fluorescence signal and contrast were achieved throughout the tissue volume to enable quantitative 3D analysis without additional tissue clearing.

A key advantage of the presented imaging protocol is its ability to visualize EHT organization across spatial scales within the same intact tissue construct. The combination of iSPIM imaging, IT-IF staining, deconvolution, and volumetric rendering enabled visualization from millimeter-scale EHT architecture to subcellular structures such as Cx43 localization. Such multiscale imaging remains challenging in intact EHTs, as conventional fluorescence microscopy is often limited in imaging depth [58,62], whereas histological approaches require physical sectioning. The same volumetric image data also enabled quantitative assessment of cellular composition throughout the tissue volume. The analysis revealed approximately 70% CM+ nuclei together with substantial regional variation in CM+ nuclei distribution, both laterally and with depth. Although EHTs were generated from a purified CM population, residual non-CMs are commonly present even after purification [63,64]. Moreover, given the limited proliferation capacity of hiPSC-CMs [65], non-CMs may increase in relative number during *in vitro* culture. While the EHTs analyzed here were generated from a CM-enriched population, the ability to quantify cellular composition and spatial heterogeneity may also prove valuable for studies of co-culture EHTs incorporating multiple cell types. The reported CM content values should be considered estimates, as classification relied on cTnT intensity thresholding and spatial association with nearby nuclei. Future membrane labeling and cell-type markers could improve the accuracy of cell identification.

Aligned CMs constitute the principal contractile architecture of the heart [13], making CM organization an important parameter for structural characterization of EHTs. In the present study, slice-by-slice orientation analysis of cTnT-labeled iSPIM 3D image stacks revealed consistent alignment along the longitudinal axis of the EHT throughout most of the tissue volume. The observed alignment is consistent with the role of passive mechanical loading provided by the two-pole EHT configuration in promoting tissue organization, in agreement with previous histological and electron microscopy findings [5]. Despite the overall high degree of alignment, significant differences were observed between tissue regions, indicating local variation in CM organization. Comparison with confocal microscopy showed visually similar tissue organization, supporting the iSPIM observation. However, the observed differences in orientation analysis may reflect optical clearing and tissue flattening following detachment for confocal imaging. The potential effects of these preparation steps on sarcomeric organization require further investigation.

The quantitative structural parameters described above demonstrate the potential of volumetric 3D analysis as a complement to existing EHT characterization methods. EHT maturation during *in vitro* culture has been associated with increased force generation, more adult-like electrophysiological properties, and improved structural organization [5]. Various methods have been developed to enhance EHT maturation, including co-culture [18,19], ECM engineering [66–68], and mechanical or electrical stimulation [69,70]. Evaluation of these approaches has largely relied on functional readouts such as force generation, beating rate, and calcium handling. In addition, efforts have been made to standardize EHTs for disease modeling and drug testing applications by applying force-based criteria prior to conducting pharmacological testing [5]. While these metrics are important indicators of tissue function, they provide limited insight into the underlying tissue organization and remodeling. The quantitative 3D structural analyses presented here provide complementary structural metrics that improve tissue characterization and contribute to future EHT standardization. Although the iSPIM imaging protocol is currently limited to end-point analysis, the structural parameters quantified here could be combined with force measurements during culture to investigate relationships between tissue organization and functional performance.

While the presented iSPIM platform provides reliable volumetric imaging and quantitative characterization of intact EHTs, the current optical configuration involves several trade-offs. In particular, the combination of a Gaussian-profile light-sheet and the requirement to image relatively large tissue volumes over a large FOV result in reduced axial resolving power. Although a thinner light-sheet could improve axial resolution, this would involve a trade-off with FOV. Alternative excitation methods such as Bessel beam or lattice light-sheet illumination could improve optical sectioning and image isotropy [30,71], but would require more complex optical instrumentation. In addition, multi-view acquisition and advanced computational reconstructions could further improve image quality and isotropy [72–75]. Signal reduction with imaging depth remains an inherent challenge in thick biological samples and is primarily caused by light scattering and optical aberrations. Future improvements can be achieved through technical developments such as adaptive optics to compensate for sample-induced aberrations [76–78], as well as through sample processing approaches [58,60]. Such strategies could further improve image quality in deeper tissue regions and quantitative characterization of EHTs. While these areas provide opportunities for future improvement, the presented single-illumination Gaussian-profile iSPIM offers a practical advantage with its relatively simple optical design. This may facilitate implementation and adaptation in other laboratories, supporting broader adoption of the approach.

Overall, the findings highlight the importance of volumetric analysis for resolving regional and depth-dependent variations in EHT organization that may not be fully captured by 2D imaging approaches. This proof-of-concept study demonstrates that the presented imaging method enables quantitative characterization of CM alignment and spatial heterogeneity while maintaining the EHT in its intact configuration.

## 5. Conclusions

The increasing complexity and physiological relevance of 3D in vitro tissue-like models have highlighted the need for imaging modalities that can provide volumetric information while preserving the models in their cultured state. In this study, we demonstrated 3D imaging and structural assessment of intact EHTs using the iSPIM platform combined with IT-IF staining and image analysis for structural quantification of EHT architecture. This approach enabled volumetric visualization of EHTs and depth-resolved structural analysis, including assessment of CM alignment and spatial registration of segmented nuclei in 3D from cTnT labeling. The results demonstrated that the overall tissue architecture well aligned with the EHT main contractile directions However, our results also revealed heterogeneity in EHTs demonstrated as regional variation in in both CM alignment and CM content, highlighting the need for more detailed structural assessment of the EHTs. Together, the presented platform and workflow are well suited for supporting studies of EHT maturation, remodeling, and disease-related structural alterations.

## Declarations

### Ethics approval and consent to participate

All experiments were performed in accordance with relevant guidelines and regulations.

### Consent for publication

Not applicable

### Availability of data and materials

The datasets used and analyzed during the current study are available from the corresponding author on reasonable request.

### Author contributions

BB, JH, and KW designed the study and concept of research. BB and KW developed and characterized the iSPIM platform. KW conducted the experimental parts and drafted the manuscript. LK differentiated the hiPSC-CMs, constructed the EHTs, and drafted the applicable parts of the manuscript. JK manufactured and provided the EHT platform. AA participated in formulating the image analysis tools. BB, AA, MP-M, JH, and LK reviewed the manuscript for finalization. All authors have read and approved the submission of this manuscript.

## Acknowledgements

The authors want to thank Jenni Roivas for the valuable segmentation work with Ilastik and summer intern Johanna Kujala for the help during building of the iSPIM setup. The authors acknowledge the Biocenter Finland and Tampere Zebrafish Laboratory for their service.

