## Additional file 1 for "Quantitative 3D imaging of intact engineered heart tissues using inverted selective plane illumination microscopy"

### Supplementary Materials and Methods

#### S1 Confocal microscopy of cleared, detached engineered heart tissues

An EHT stained with cTnT, Cx43 and nuclei was first imaged as intact tissue between the poles with the iSPIM platform. Then, the EHT was removed from the poles with forceps and cleared with RapiClear 1.49 (SunJin Lab Co., Taiwan) tissue clearing reagent following the manufacturer’s protocol. The cleared EHT was then imaged with Evident FV4000 confocal laser-scanning microscope (Evident, Japan) using a 20X magnification, 1024x1024 pixels and acquiring the full tissue Z-stack with step size of 1 µm. After acquiring, matching cellular structures were found from both iSPIM and confocal images, and the same CM alignment analysis was performed to both imaging modalities as described in section 2.13. Prior to the analysis, the confocal image stack was rotated to match vertical tissue orientation in Fiji.

### Supplementary Results

#### S1 3D alignment of cardiomyocytes in detached and cleared EHTs

CM alignment was quantified both from intact iSPIM imaging and detached, cleared tissue confocal microscopy images to assess the changes occurring during further sample handling. Similar sections of the same tissue were found from both imaging modalities (Fig. Sa). The quantification of dominant angle orientation showed significant differences in angle distribution and mean orientation angle (LSCM: 87.7° [86.1-88.9] vs. iSPIM: 89.7° [89.1-91.1]), indicating changes in the tissue structure caused by sample detachment, tissue clearing protocol, and different mounting for confocal imaging (Fig. Sb). These results demonstrate the advantage of iSPIM for imaging intact tissues, still in their cultured configuration, resulting in the most accurate analysis of tissue architecture.

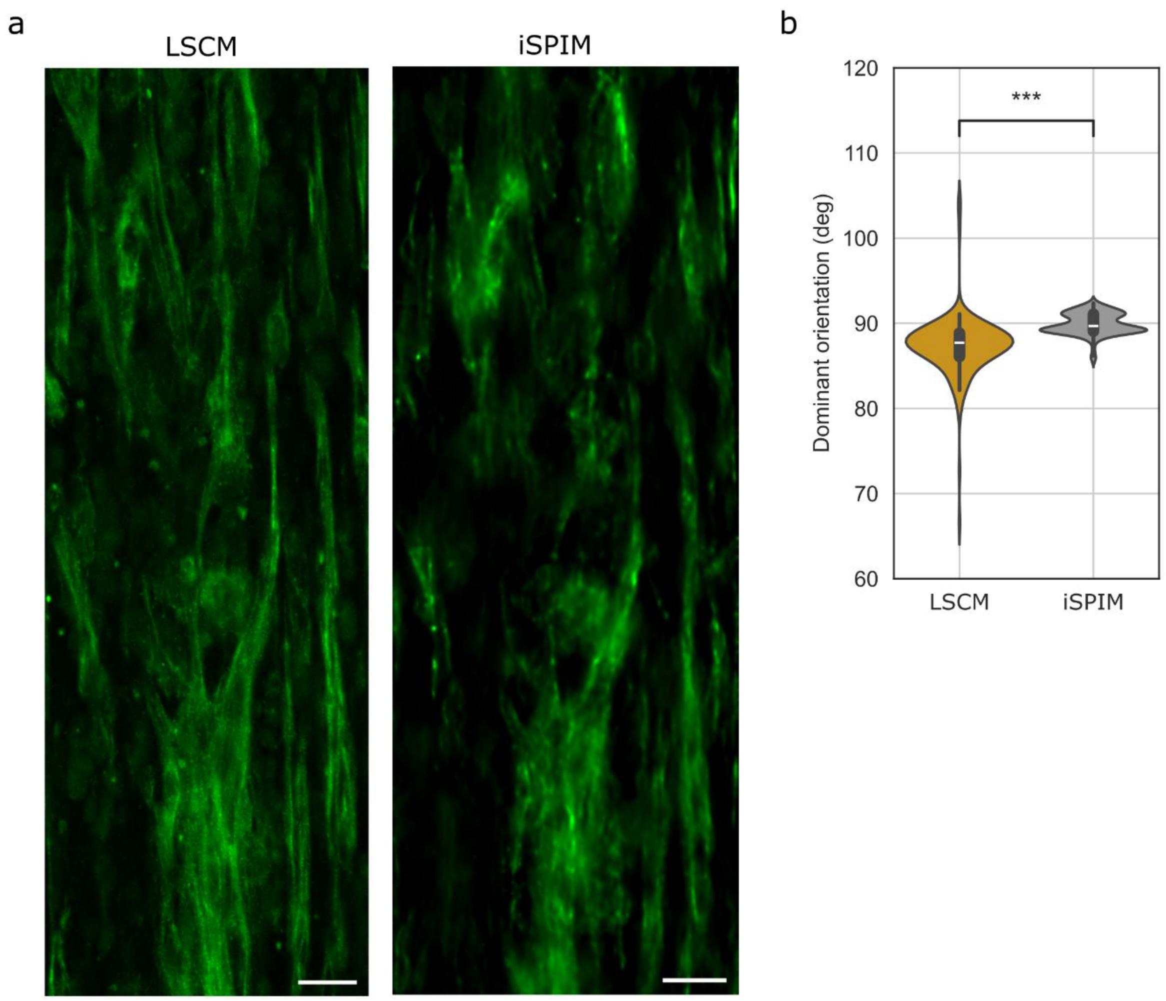


**Fig. S Comparison of alignment with intact iSPIM imaging and detached confocal imaging**. **a** 7 µm thick maximum intensity projection (MIP) of similar engineered heart tissue (EHT) area labeled with cardiac-specific troponin T (cTnT), imaged with laser scanning confocal microscope (LSCM; left) and inverted selective plane illumination microscope (iSPIM; right). Scalebar: 25 µm. **b** Dominant cardiomyocyte orientation measured from the cTnT labeled volumes. Statistical significance: ***$p<0.001$. $n=169$ for LSCM and $n=380$ for iSPIM

Additional files

Additional Video S1. Representative intact EHT volumetric visualization with cardiac-specific troponin T visualized in green and nuclei in blue. Duration: 33 s.
